# Real Time *de novo* Deposition of Centromeric Histone-associated Proteins Using the Auxin Inducible Degradation system

**DOI:** 10.1101/304725

**Authors:** Sebastian Hoffmann, Daniele Fachinetti

## Abstract

Measuring protein dynamics is essential to uncover protein function and to understand the formation of large protein complexes such as centromeres. Recently, genome engineering in human cells has improved our ability to study the function of endogenous proteins. By combining genome editing techniques with the Auxin Inducible Degradation (AID) system, we created a versatile tool to study protein dynamics. This system allows us to analyze both protein function and dynamics by enabling rapid protein depletion and re-expression in the same experimental set-up. Here, we focus on the dynamics of the centromeric histone-associated protein CENP-C, responsible for the formation of the kinetochore complex. Following rapid removal and re-activation of a fluorescent version of CENP-C by auxin treatment and removal, we could follow CENP-C *de novo* deposition at centromeric regions during different stages of the cell cycle. In conclusion, the auxin degradation system is a powerful tool to assess and quantify protein dynamics in real time.

## 1. INTRODUCTION

The ability to follow protein dynamics in vivo is extremely important to determine protein function and the role of specific proteins in complexes. This is particularly important for the assembly of large protein complexes, in which the association of specific proteins may rely on the presence of others. Therefore, the analysis of the dynamics of each complex component may yield essential information on the global function and the assembly of the complex itself.

Centromeres are DNA/protein structures necessary to maintain the balance in genetic information by controlling faithful chromosome segregation during cell division. They are the foundation for the assembly of the kinetochore that, in turn, is required to interact with spindle microtubules. Centromeres are epigenetically identified by the presence of a histone H3 variant named CENP-A. CENP-A is strongly enriched at centromeric regions and is the base for the assembly of the entire centromere/kinetochore complex. Altogether, the centromere/kinetochore can form a complex with over 100 proteins of about 150 nm in human cells [1]. Recently it was proposed that the subunits of the Constitutive Centromere Associated Network (CCAN) interact between each other in an interdependent manner, and that this interdependency is necessary for the stability of the entire complex [2, 3]. Although most of the centromeric proteins are present along all stages of the cell cycle, their assembly is highly regulated and often restricted to a limited time window [4]. For example, CENP-A is deposited only once every cell cycle, at the exit of mitosis [5] via a tight regulatory mechanism [6].

Here we describe the <u>A</u>uxin <u>I</u>unducible <u>D</u>egradation (AID) system as a method to track protein dynamics throughout the cell cycle. The AID system is based on the transplantation in eukaryotes [7] of a ligand-induced degradation system found in plants [8, 9]. Addition of the plant hormone indole-3-acetic acid (IAA, auxin) mediates the interaction of the AID-tagged protein of interest with the ectopically expressed F-box from plant (Transport Inhibitor Response 1; TIR1), that can associate with Skp1 in eukaryotic cells and form an SCF-TIR1 complex to induce protein ubiquitination and degradation. AID size is about 25 kDa (similar to GFP), but can be reduced by ∼1/3 without loss of functionality, as observed in budding yeast [10].

The auxin degradation system has already been successfully used in the past [2, 11–15] to study protein function following rapid and complete protein degradation at every cell cycle stage. As an example, using this system we recently demonstrated that CENP-A is dispensable for maintenance of an already assembled centromere/kinetochore complex [11].

In this chapter, we illustrate how to exploit one of the key characteristics of the AID system, its reversibility. We describe in detail how to (Figure 1A): i) generate a stable human cell line expressing the E3 ubiquitin ligase TIR1, ii) insert an AID tag coupled with a fluorescent protein (mRFP or EYFP) at an endogenous gene locus, iii) select and screen for correct AID integration and iv) measure protein dynamics by time lapse microscopy following rapid removal and reactivation of the AID-tagged protein pool (Figure 2). Here, we focus on the histone CENP-A-associated protein CENP-C, the major component for kinetochore assembly in human cells. However, in principle, any protein of interest can be tagged and assayed for protein dynamics including histones, as we have previously shown for CENP-A [11].

**Figure 1:**
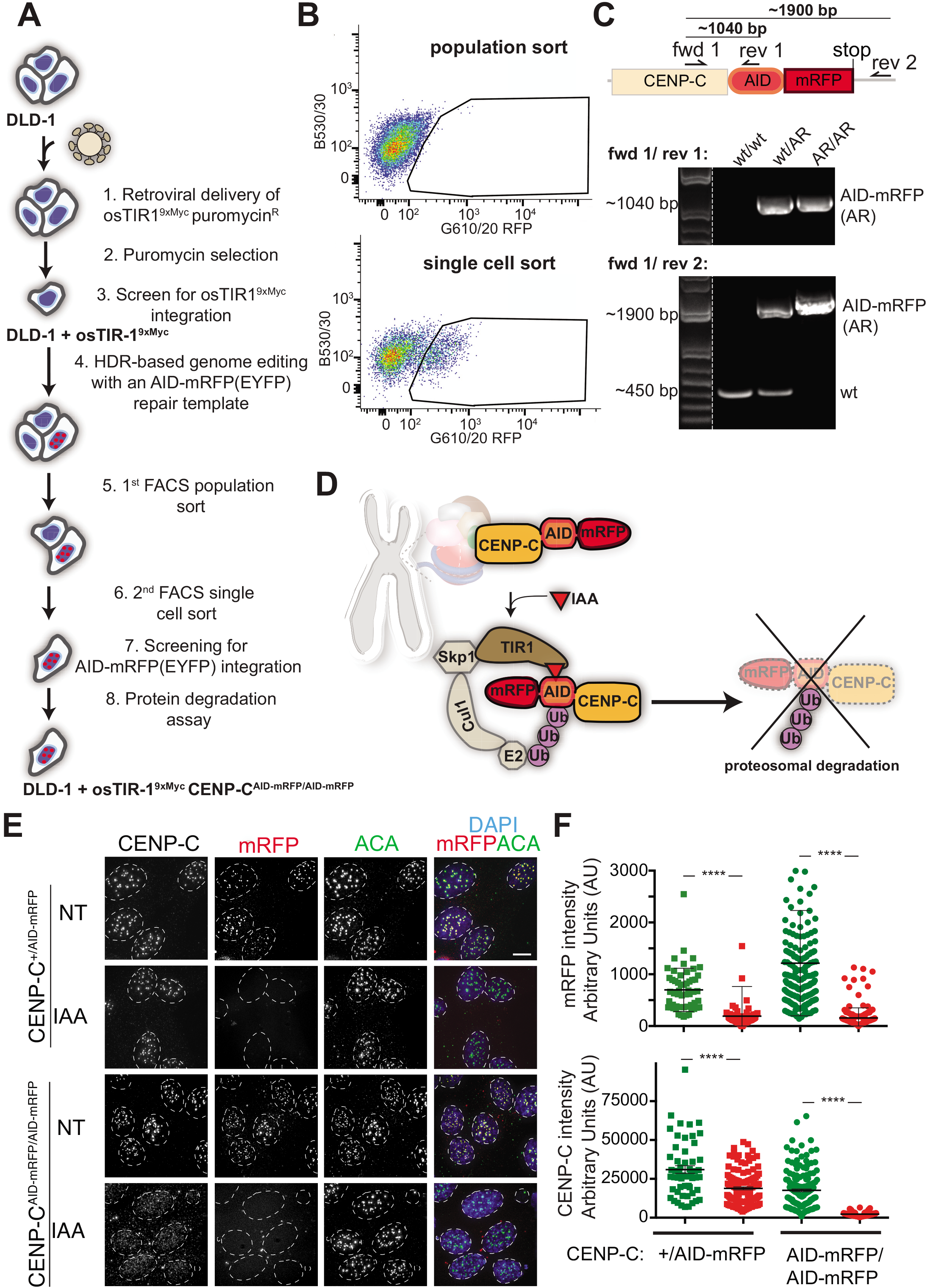
CENP-C genome engineering. **A**. Schematic of the TALEN-mediated genome editing strategy to endogenously tag CENP-C with AID and mRFP (or EYFP) in cells expressing osTIR1. **B.** A two-step FACS selection procedure of cells in which CENP-C has been endogenously tagged with an AID and mRFP. In step one (upper plot) a population of red fluorescent cells was collected. In step two (lower plot) single red fluorescent cells were sorted into a 96-well plate. **C.** CENP-C genotypes validated in the indicated cell lines using PCR to distinguish normal (+/+), single tagged (+/AID-mRFP) or double tagged (AID-mRFP/ AID-mRFP) cells **D.** Schematic illustration of the AID system to induce CENP-C degradation including the one incorporated at centromeric regions that interact with the CENP-A nucleosome (in red) and the CCAN complex (multicolor). **E.** Representative immunofluorescence images of cells showing CENP-C depletion after 24-hours treatment with IAA. ACA was used to mark centromere position. Scale bar = 5 μm. **F.** Quantifications of the experiment in E. Dots represent single centromere. Unpaired t test: **** p < 0.0001.

**Figure 2.**
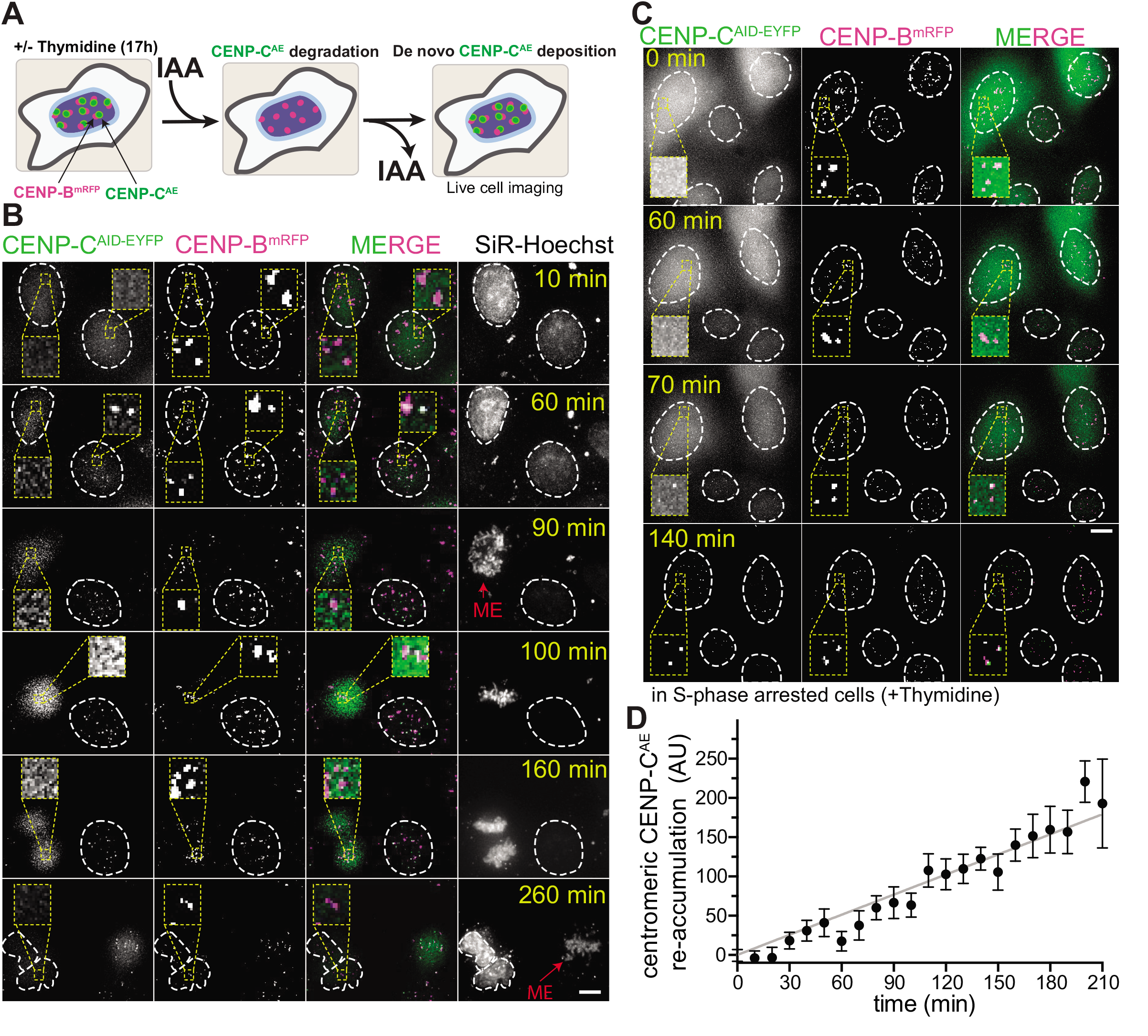
Real time *de novo* deposition of CENP-C following rapid protein deletion and reactivation. **A.** Experimental scheme to follow *de novo* deposition of CENP-C^AID-EYFP^ in a DLD-1 cell line. **B-C.** Representative live cell images of CENP-C^AID-EYFP^ reloading in asynchronous DLD-1 cells (B) or in S-phase arrested cells (C). The inset highlighted with a yellow dashed square shows a magnification of a section of CENP-B-marked centromeres. Red arrows mark cell entry into mitosis (Mitotic Entry, ME). Images were acquired 1h after IAA wash-out. **D.** Graphs represent the mean from 10 cells of CENP-C re-expression and re-accumulation at centromeric regions marked by CENP-B-mRFP in S-phase arrested cells followed by live cell imaging. Error bars represent the SEM. AU = Arbitrary units. Scale bar = 5 μm.

## 2 MATERIALS

### 2.1 Cell culture reagents

1. DMEM-GlutaMAX (Gibco or equivalent) supplemented with 10% fetal bovine serum (GE Healthcare), 100 U/ml penicillin, 100 U/ml streptomycin
3. Trypsin (Gibco or equivalent)
4. Phosphate-buffered saline (PBS): 137 mM NaCl, 2.7 mM KCl, 10 mM Na_2_HPO_4_, 1.8 mM KH_2_PO_4_
2. Cell culture incubator (5% CO_2_, 37°C)
3. 500 mM Auxin Indole-3-Acetic Acid sodium salt (IAA) in water. Use at 500 μM final concentration.
4. 10 mg/ml Puromycin. Use at 2 μg/ml final concentration
5. 10 mg/ml Polybrene. Use at 8 μg/ml final concentration
6. 100 mM Thymidine dissolved in water. Use at 2 mM final concentration

### 2.2 Cell line generation

1. FuGENE HD (Promega) (or equivalent transfection system)
2. VSV-G plasmid
3. Opti-MEM Reduced Serum Media (Gibco)
4. Cell Line Nucleofector Kit V, Lonza Nucleofector Device
5. QIAGEN Plasmid Midi Kit (or equivalent plasmid preparation system)
6. SURVEYOR Mutation Detection Kit (Transgenomic or equivalent)

### 2.3 Cell selection

1. Fluorescence activated cell sorting (FACS) device.
2. FACS buffer: 1% BSA, 5 mM EDTA, 5 mM HEPES in 1x PBS

### 2.4 Cell screening

#### 2.4.1 PCR

1. QuickExtract™ DNA Extraction Solution (Epicentre)
2. NanoDrop
3. Q5 Hot Start High-Fidelity 2X Master Mix (NEB)
4. Oligomers flanking AID-mRFP at the C terminus of CENP-C:

> 5’-GTTAGAGGAATCCACAGCAGT-3’ (forward)
>
> 5’-TTACAAAGACAAATATTCCAACTA-3’ (reverse)
>
> 5’-CTCATGAAAGGATCGGATGC-3’ (reverse, binds in the AID sequence)
5. Thermocycler
6. Agarose gel electrophoresis system
7. 1% agarose gels (w/v)
8. GelGreen stain (Interchim or equivalent)

#### 2.4.2 Immunofluorescence microscopy

1. Triton blocking buffer: 0.2 M Glycine, 2.5% FBS, 0.1% triton X-100 in 1x PBS.
2. Primary antibodies: CENP-C (1:1,000 MBL International, guinea pig), CENP-B (1:1,000 Abcam, rabbit)
3. Secondary antibodies: Fluorophore conjugated anti-guinea pig and anti-rabbit antibodies.
4. DAPI
5. Anti-fading mounting reagent (Life technologies or equivalent)

#### 2.4.3 Immunoblot

1. Sample buffer: 62.5 mM Tris, pH 6.8, 2.5% SDS (w/v), 10% Glycerol (v/v), 5% β-Mercaptoethanol (v/v), 0.002% bromophenol blue (v/v)
2. Running buffer: 25 mM Tris Base, 192 mM Glycine, 0.1% SDS (w/v)
3. Blocking solution: 5% milk (w/v) in TBS with Tween (0.1%)
4. 4-15% Mini-PROTEAN TGX Precast Protein Gel (Bio-Rad)
5. Trans-Blot Turbo Transfer System (Bio-Rad)
6. Trans-Blot Turbo Midi Nitrocellulose Transfer Packs (Bio-Rad)
7. Tris-buffered saline (TBS)-tween: 50 mM Tris-HCl, pH 7.5, 150 mM NaCl, 0.1% tween (v/v)
8. Antibodies for immunoblot: Anti-Myc-tag, clone 4A6 (Merck chemicals, ref.: 05-724, used at 1 μg/ml in blocking solution), ECL™ Anti-mouse IgG, Horseradish Peroxidase linked whole antibody (from sheep, GE Healthcare UK, used 1:10,000 in blocking solution)
9. Pierce HCL (Thermo Scientific or equivalent)
10. Gel Imaging System

### 2.5 Live cell imaging

1. μ slide 8-well (IBIDI Cat. No. 80826 or equivalent).
2. Deltavision core system equipped with a CoolSNAP_HQ2 Camera and an Olympus UPlanSApo 100x oil-immersion objective (numerical aperture 1.4) for fixed cells and an Olympus 40x oil-immersion objective for live cell imaging (Applied Precision)
3. Software for data analysis: softWoRx (Applied Precision), ImageJ (open source), MetaMorph
4. CO_2_-independent medium (Gibco) supplemented with 10/ fetal bovine serum (GE Healthcare) 100 U/ml penicillin, 100 U/ml streptomycin
5. SiR-DNA dye kit (Spirochrome)

## 3. METHODS

### 3.1 Cell line generation

#### 3.1.1 Retroviral mediated osTIR1 transgene integration

Auxin mediated degradation of an AID-tagged protein is only achievable in the presence of the plant E3 ubiquitin ligase TIR1 (transport inhibitor response 1). To stably express TIR1 trans-gene from rice *(Oryza sativa; os*TIR1) into human cells (in our case colorectal adenocarcinoma DLD-1) this procedure exploits retrovirus-mediated integration. There are several methods to deliver DNA to mammalian cells in tissue culture. Here, we describe a method using Fugene HD (Promega) transfection reagent.

##### Retrovirus production

1. Day 0: Seed 3×10^6^ 239-GP cells in a 10-cm cell culture plate (total volume: 10 ml).
2. Day 1: Transfection. Add 15 μl Fugene HD to 500 μl OptiMEM or free FBS/antibiotics DMEM medium and vortex 1s (use pre-warmed medium).
3. Add 3 μg VSV-G and 5 μg osTir1-9xMyc construct in a pBABE vector backbone comprising a puromycin resistance cassette.
4. Incubate transfection mix for 15 min at RT and add the mix dropwise onto the cells (biosafety measures for working with the retroviruses need to be adopted at this point)
5. Day 2: Replace culture medium with 6 mL fresh medium. Allow virus production for another 48 h.
6. Day 4: Collect 6 mL virus-containing medium and filter the medium through a 0.45 μm filter.
7. Aliquot the virus, snap freeze in liquid nitrogen and store at −80°C or use directly (see below).

##### Retrovirus infection

Bio-safety measures for working with the virus are required

8. Day 0: seed DLD-1 cells in a 6-well cell culture plate at 20% confluency (3 wells/transfection).
9. Day 1: replace medium with 2 mL of culture medium containing 8 μg/ml Polybrene.
10. Day 2: add 250 μl, 500 μl and 750 μl of the virus to three separate wells and allow virus infection for 2 days.
11. Day 3: Wash-out the virus three times with 2 mL culture medium. Wash cells once with 2 mL 1x PBS, remove PBS and add 300 μl trypsin to detach cells from the tissue culture dish. Incubate cells in trypsin for 5 min in the incubator. After about 2-5 min shake of the cells. Confirm cell detachment using a bright light microscope. When all cells are detached from the plate re-suspend cells in 1.7 ml culture media. Pool and seed infected cells in a 15-cm cell culture plate. Add culture medium (total volume: 25 ml).
12. Day 4: add puromycin to select for cells with osTIR1 insertion (final puromycin concentration: 2 μg/ml for the DLD-1 cell line).

##### osTIR1 screening

Two to seven days after puromycin addition refresh the cell culture medium in order to remove dead cells and to ensure puromycin efficacy. Two weeks after puromycin addition it is possible to observe the formation of puromycin resistant colonies. For colony isolation, proceed as follows:

13. Sterilize small (˜ 1cm^2^) Whatman paper pieces.
14. Soak the paper in trypsin. Remove the excess of trypsin by squeezing the paper against an empty sterile surface (e.g. cell culture plate).
15. Wash the puromycin resistant colonies with 15 ml 1x PBS. Remove PBS.
16. Place the Whatman paper from step 14 directly onto a colony using sterile forceps.
17. After around 5 min in the cell culture incubator, cells start to detach. At this point, convey the colony into a 24-well plate containing 0.5 ml pre-warmed cell culture medium by transferring the whole paper.
18. On the following day, remove the Whatman paper and replace the medium.

##### Immunoblot

Next, use immunoblot using an anti-Myc antibody to screen all single cell-derived colonies for genomic integration and expression of osTIR1-9x-Myc construct.

19. Once the single cell clones are confluent collect cells by trypsinization as described in step 11. Here, use 100 μl of trypsin for each well of a 24-well plate and re-suspend cells with 400 μl culture media. Seed 100 μl of cells into a 12-well plate in duplicate. Add culture media to the 12-well plate (total volume: 1 ml). Use the remaining 400 ul for western blot analysis.
20. Collect the 400 μl cells in a 1.5 ml Eppendorf tube. Harvest cells by centrifugation (2000 g for 5 min).
21. Carefully remove the supernatant and wash the cell pellet once with 1x PBS. Re-suspend cells in 150 μl sample buffer.
22. Boil samples at 100°C for 10 min and separate proteins of the cell extracts by SDS-polyacrylamide gel electrophoresis (PAGE). Use a 10/ Mini-PROTEAN^®^ TGX™ precast protein gel and a biorad electrophoresis system or similar. Insert a SDS-PAGE gel into the designated electrophoresis system and add running buffer to the system. Load 30 μl protein samples and 10 μl of a protein ladder in separate gel pockets using a Hamilton syringe or gel loading tips for pipettes. Apply low voltage (70 V) for accurate protein separation.
23. After electrophoresis transfer proteins onto a nitrocellulose membrane with the Biorad Trans-Blot^®^ Turbo™ Transfer System or equivalent.
24. After transfer, block the membrane in 25 ml blocking solution for 30 min at room temperature in a box that allows full coverage of the membrane.
25. Incubate the membrane with 10 ml primary antibody against the myc-tag in 0.5/ milk TBS-Tween for 2 h at room temperature. Ensure that the membrane is completely covered.
26. Wash the membrane 3X with 25 ml TBS-Tween at room temperature (5 min per wash).
27. Incubate 45 min with 10 ml secondary antibody linked to horseradish peroxidase at room temperature.
28. Wash 3X for 10 min with 25 ml TBS-Tween at room temperature.
29. Develop the membrane using Pierce HCL and a gel imaging system.

Based on the western blot result, pool single cell-derived colonies with similar high expression levels of osTIR1-9x-Myc (Note 1).

Use this cell population for the subsequent genome editing process as described in the following sections.

#### 3.1.2 AID-tag integration at the CENP-C locus

In the next step, introduce an AID tag combined with a fluorescent protein [here: monomeric red fluorescent protein (mRFP) or enhanced yellow fluorescent protein (EYFP)] at the endogenous loci of a centromeric histone-associated protein (in this case CENP-C, but a similar strategy was used to target the histone H3 variant CENP-A). To this end, this procedure describes use of a TALEN genome editing strategy (<u>T</u>ranscription <u>A</u>ctivator-<u>L</u>ike <u>E</u>ffector binding domain coupled with a FokI <u>N</u>uclease), however, it can easily be adapted for a CRISPR/Cas9 or any other genome editing strategy (Note 2). The AID-mRFP(EYFP)-tag is integrated at the target site via the homology directed repair (HDR) (Note 3).

In this case, genome targeting plasmids were delivered by electroporation using a Lonza Cell Line Nucleofector Kit and the Lonza Nucleofector electroporation device (but other transfection methods could be used).

##### Electroporation

1. One day before transfection, grow DLD-1 cells in a 10-cm cell culture dish in antibiotics-free cell culture medium (total volume: 10 ml).
2. Prepare a 1 μg plasmid mix containing 800 ng repair template plasmid, 100 ng left TALEN plasmid and 100 ng right TALEN plasmid in a 1.5 ml Eppendorf tube (use Midi-Prep DNA quality for all plasmids). In addition to the transfection of the gene targeting plasmids we also recommend to perform a separate control by transfecting 500 ng of a plasmid encoding soluble GFP.
3. Trypsinize cells as described previously (here: use 1.5 ml trypsin and re-suspend cells with 8.5 ml culture medium). Collect cells in a 15 ml Falcon tube. From a fully confluent 10-cm cell culture dish 3 transfections can be performed.
4. Centrifuge cells for 5 min at 1000 g at room temperature.
5. Aspirate medium completely and re-suspend cells in 100 μl (for each transfection) of Lonza Cell Line Nucleofector kit V buffer with supplement (Note 4).
6. Add re-suspended cells to the plasmid cocktail and mix gently by pipetting up and down. Wait 5 min.
7. Transfer the plasmid/cell mixture without generating bubbles into an electroporation cuvette provided in the nucleofector kit.
8. Place the cuvette in the Lonza Nucleofector device and apply program U-017 (Note 4).
9. Transfer cells carefully with a plastic pipette (provided in the kit) to a 6-well cell culture plate containing 2 ml pre-warmed cell culture medium without antibiotics. Shake the plate gently to dispense cells evenly.
10. One day post-transfection refresh the medium supplemented with antibiotics. Check efficiency of transfection using a fluorescence microscope (e.g EVOS FL Cell Imaging System). Transfection efficiencies of 70% to 90% with 50-90% cell viability are commonly reached.
11. Grow cells for around three to five days and then expand to obtain a bigger number of cells.

##### Clones Selection

In order to select for DLD-1 clones in which the AID-mRFP/EYFP tag has been inserted a two-step FACS selection protocol was used (Note 5).

12. Three to five days after transfection, harvest cells in 15 mL Falcon tube by trypsinization as described above.
13. Wash cells with 5 ml FACS buffer and re-suspend cells in 0,5 to 1 ml FACS buffer. Also prepare a negative control with non-transfected DLD-1 cells in the same way.
14. Sort mRFP positive cells in a tube and plate cells in a cell culture dish (figure 1 B, upper panel) (Notes 6, 7).
15. Once cells are happily growing and a good confluency is achieved (normally after 2-5 days), perform a single-cell sorting into a 96-well plate using filtered conditional medium (50% fresh medium, 50% 3 days old medium) (see Note 8). At this time, a strong enrichment of RFP positive cells can be observed (figure 1 B, lower panel).

Before performing single cells sorting, screening of cells for positive integration using PCR screening is recommended (see below).

#### 3.1.3 Clones Screening

Following FACS selection there are multiple possibilities to screen for AID-mRFP/EYFP integration.

Here we will present three of them (Note 9).

##### Cell viability screening

The most straightforward way is to initially screen for cell viability if the protein of interest is essential as in the case of CENP-C. To this end:

16. Seed clones derived from single cell sorting into duplicate 24-well plates containing regular medium or regular medium + IAA (total volume: 0.5 ml).
17. Clones that die within 4 days of IAA treatment are selected.
18. Expand the corresponding untreated single cell clones.
19. Confirm the integration of AID-mRFP in these clones by PCR (as described below) and immunofluorescence microscopy.

##### PCR examination for AID-mRFP integration

Perform standard PCR to test for the integration of the AID-mRFP (EYFP) tag (Note 10). DNA extraction using QuickExtract™ DNA Extraction Solution:

20. Collect cells from a 1 mL fully confluent 12-cm cell culture dish by trypsinization and centrifugation in a 1.5 mL Eppendorf tube.
21. Wash cell pellets with PBS and re-suspend in 50 μl QuickExtract™ DNA Extraction Solution (scaling of cell pellet/solution is possible) (Other DNA extraction procedures can be used).
22. Incubate 5 min at RT.
23. Vortex the DNA extraction mix for 10 s.
24. Incubate 6 min at 65°C.
25. Vortex the DNA extraction mix for 15 s.
26. Incubate 2 min at 98°C.
27. Vortex the DNA extraction mix for 15 s.
28. Cool down DNA for PCR reaction (store at −20°C).
29. Measure DNA concentration using a NanoDrop™ (use 300 ng/μl final DNA concentration for the PCR).
30. Prepare the PCR mix according to the table below.
31. Run PCR reaction according to the settings below.

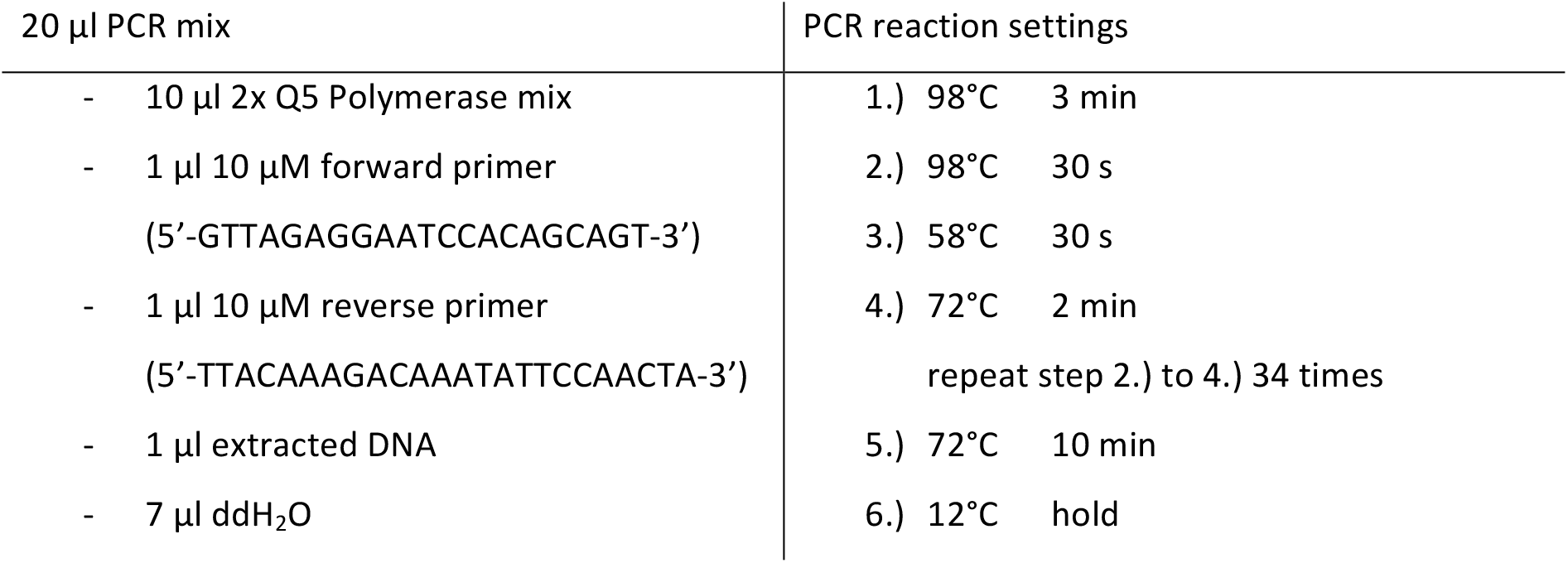

32. Analyze the PCR results by gel electrophoresis using a 1% (w/v) agarose gel containing GelGreen.
33. Excise the PCR products from the gel.
34. Purify the PCR product by using a standard DNA-agarose gel extraction kit and use a Sanger sequencing service to bi-directionally sequence the PCR product.
35. Verify the successful integration of the AID-mRFP/EYFP tag at the CENP-C locus with the sequencing results.

##### Immunofluorescence microscopy screen

CENP-C^AID-mRFP/AID-mRFP^ clones identified by the viability screening and PCR analysis can be seeded for immunofluorescence microscopy analysis in duplicate on coverslips in a 24-well plate.

36. In order to test for the degradation of CENP-C^AID-mRFP^ add IAA overnight to one of the slides prior to fixation (Figure 1 D, E) (Note 11).
37. Pre-extract cells with 0.5 ml 1x PBS-T (0.1% Triton X-100) for 1 min at room temperature.
38. Fix cells by adding 200 μl 4% formaldehyde for 5-10 min at room temperature.
39. Wash out Formaldehyde with 0.5 ml 1x PBS-T (2 times).
40. Incubate 30 min in 0.5 ml triton blocking buffer at room temperature.
41. Incubate with 150 μl primary antibodies in triton blocking buffer for 1 h (anti-CENP-C and anti-CENP-B) at room temperature.
42. Wash 3x with 0.5 ml PBS-T.
43. Incubate with 150 μl secondary antibodies in triton blocking buffer for 45 min at room temperature.
44. Wash 3x with 0.5 ml PBS-T.
45. Incubate with 150 μl DAPI in PBS (1 μg/ml) in PBS for 10 min at room temperature.
46. Mount coverslips with anti-fading reagent.

Acquire fluorescence microscopy images using a fluorescent microscope such as the DeltaVision Core system (Applied Precision). We use a 100x Olympus UPlanSApo 100 oil-immersion objective (numerical aperture 1.4) and a 250W Xenon light source. The system is equipped with a Photometrics CoolSNAP_HQ2 Camera. Acquire 4 μm z-stacks (step size: 0.2 μm). Quantification of CENP-C using an automated system [16] and ACA to mark centromeres reveals that CENP-C is completely depleted at the centromere after addition of IAA (Figure 1 F).

### 3.2 Auxin mediated depletion, re-expression and reloading of CENP-C^AID-EYFP^ at the centromere

The auxin degradation system is a useful tool to study consequences of rapid and complete protein depletion. Due to its reversibility, the system can also be used to study *de novo* re-expression and localization of AID-tagged proteins including histone variants. Here we use the auxin degradation system to follow *de novo* deposition of CENP-C following its complete degradation. To study the dynamics of CENP-C deposition along the cell cycle we used the cenP-C^AID-EYFP/AID-EYFP^ cell line in which also one allele of CENP-B has been endogenously tagged with an mRFP-tag (the CENP-B tagging was carried out in a similar way as described in the previous section). The presence of mRFP on CENP-B allows us to mark centromere position and to follow *de novo* CENP-C deposition by live cell imaging. In order to completely deplete CENP-C^AID-EYFP^ (henceforth referred to as CENP-C^AE^) we added IAA to the culture medium. After CENP-C^AE^ depletion we washed-out auxin and allowed the re-expression of CENP-C^AID-EYFP^. We followed CENP-C^AE^ reloading at the centromere by live-cell imaging in asynchronous cells or in cells arrested in S-phase (Figure 2 A).

#### 3.2.1 Cell preparation

1. Day 0: seed cells in 5 adjacent wells of an 8-well IBIDI slide (total volume: 300 μl).
2. Day 1: replace media with 300 μl 2 mM thymidine-containing cell culture medium overnight for the S-phase arrested condition.
3. Day 2: replace media with 200 μl IAA (500 μM final concentration) for 6 h (except for the Not Treated – NT condition) and also maintain thymidine for S-phase arrested cells.
4. Wash out IAA carefully three times with 300 μl culture medium.
5. Leave cells for 15 min in tissue culture incubator and repeat washes (Note 12). During this time incubate cells with SiR-DNA dye kit (working solution 1 μM).
6. Replace media with CO_2_ independent medium and start live-cell imaging immediately (for the S-phase arrested condition maintain thymidine in the cell culture medium).

#### 3.2.2 Live cell imaging with fluorescence microscopy

7. Prior to image acquisition adjust temperature to 37°C.
8. For imaging, we use the DeltaVision Core system (Applied Precision) with an Olympus 60X/1.42, Plan Apo N Objective. The microscope is equipped with a CoolSNAP_HQ2 camera.
9. Acquire 20 μm z-stacks (2 μm step size) with a 2X2 binning. Acquire images every 10 min (Note 13).

#### 3.2.3 Experimental analysis

10. CENP-C^AE^ reloading can be qualitatively analyzed in different cell cycle phases using ImageJ (open source image processing software) after generating maximum intensity projections of the deconvolved z-stacks (Note 14).
11. To quantify CENP-C^AE^ re-expression at the centromeres over time, save un-deconvolved 2D maximum intensity projections as un-scaled 16-bit TIFF images and signal intensities determined using MetaMorph (Molecular Devices).
12. Draw a 10 × 10 pixel circle around a centromere (marked by CENP-B^mRFP^) and draw an identical circle adjacent to the centromere (background). The integrated signal intensity of each individual centromere is calculated by subtracting the fluorescence intensity of the background from the intensity of the adjacent centromere.
13. Average 10 centromeres to provide the average fluorescence intensity for each individual cell and quantify about 10 cells per experiment in S-phase arrested cells conditions.

##### Conclusion

Several techniques exist to analyze protein dynamics such as <u>F</u>luorescence <u>R</u>ecovery <u>A</u>fter Photobleaching (FRAP), photo-activation of proteins, <u>R</u>ecombination <u>I</u>nduced <u>T</u>ag <u>E</u>xchange (RITE) and SNAP-based pulse labeling (for a detailed list see table 1 in [17]). The AID system described here has several advantages over some of the aforementioned methods to analyze protein turn over. First and most important, it allows the analysis of both protein function and dynamics in the same experimental setting by inducing rapid protein depletion (by IAA) and by achieving rapid protein reexpression (by IAA removal), both processes within minutes. Auxin is permeable (it passes through the cell membranes) and easy to wash out since it is dissolved in water (no need of DMSO), and it does not require the presence of any additional components (e.g. CRE recombinase in the RITE) to start monitoring protein dynamics. Since specific IAA-mediated protein degradation does not affect mRNA, the AID-tagged protein re-accumulates very rapidly, allowing live measurement of protein turnover at short timescales in every cell (in large numbers) and at every complex (in this case centromeres). Also, it does not require any particular laser or specialized equipment, so a standard fluorescence microscope can be used.

One disadvantage of the AID technique is that an *“ad hoc”* system is required for every protein of interest, involving extensive gene modification such as gene tagging that may disrupt protein function and the insertion of a transgene (TIR1). Also, target protein dynamics completely depend on protein expression levels, and there could be some photo-bleaching events during time-lapse experiments. However, at least one of these caveats is also found in all of the techniques mentioned above, highlighting again the unique advantage of using the AID system.

In summary, here we presented the auxin degradation system as a unique tool to study the *de novo* deposition of centromeric proteins. This tool could be adapted to study a wide range of other proteins and protein complexes to gain better insight into their function and dynamics.

## 4 NOTES

1. High level of osTIR1 expression is essential for rapid and complete degradation of AID-tagged target proteins. The majority of clones express the osTIR1-9x-Myc transgene due to the puromycin selection after virus infection, however expression levels might change due to the different integration sites within the genome. In order to decide which clones to pick, band quantification on the western blot of osTIR1-9x-Myc expression level is critical. The usage of a positive reference for TIR1 expression level [13] is recommended. We commonly pool all clones that display expression levels that are at least half as strong as the level of the reference clone. An alternative strategy is to AID-tag the gene of interest before adding the TIR1. This strategy will help if different levels of TIR1 are required to degrade the protein of interest (e.g. too much TIR1 might lead to leaky protein degradation in the absence of IAA), however it will preclude the usage of the “<u>Cell viability screening</u>” method described in this book chapter.
2. We used a TALEN-based genome editing approach, which allows site-specific genomic modifications with a very low chance of off-target genome editing effects. The design of left and right TALEN DNA recognition FokI fusion constructs have been extensively described [18] and, due to space limitations, will not be addressed here. Briefly, TALENS are cloned into a modified version of pcDNA3.1 (purchased from Invitrogen) containing also a cDNA sequence for the Fok I endonuclease domain. TALENs CENP-C target sequences: GAGGAAAGTGTTCTTC and GGTTGATCTTTCATC [16]. TALENs cleavage efficiency is tested by using a surveyor assays as described in Ran et al. (2013) [19].
3. To exploit HDR in cells, a repair template plasmid (containing the *AID-mRFP or AID-EYFP* sequence flanked by a 0.3-1 kb homologous DNA sequence on each site) is co-transfected along with plasmids expressing left and right TALEN DNA recognition proteins. The homologous DNA sequences are designed up- and down-stream of the STOP genomic target site. Since HDR is a rare event we increased the repair template plasmid concentration relative to the left and right TALEN plasmids in the co-transfection mixture. Since the N-terminus of CENP-C is important for the interaction with Mis12, a crucial component for kinetochore formation [20], we decided to introduce the AID-mRFP tag at the C terminus. Previously, we have also introduced the AID-tag at the N-terminus of other proteins such as CENP-A with a very similar approach. In our opinion, the amino terminal tagging might be advantageous since following double strand break formation by TALENs or Cas9, DNA will be more frequently repaired via the non-homologous end-joining pathway. This error-prone repair can lead to rearrangements (deletion or insertion) and then, possibly, to de-activation of one of the alleles. In this case, the necessity to introduce the AID tag via HDR is reduced to one instead of two alleles.
4. The choice of the kit and the electroporation program are cell line dependent and need to be tested. Lonza provides a database for the correct choice of the kit and electroporation program but some troubleshooting might be required.
5. A one-step single cell sorting procedure after transfection commonly results in a high number of false positive cells. This is due to low AID-mRFP/EYFP integration efficiency and low fluorescence intensity of cells expressing mRFP (or EYFP). This will highly depend on the endogenous promoter of the tagged gene of interest. We first enriched for a population of mRFP positive cells. Next, after expansion of the sorted population, we performed single cell sorting for clone selection of integration of the desired tag.
6. Population sorting requires up to 1 hour since the number of mRFP (EYFP)-positive cells might be very low (˜0.1%). We sort cells in 200 μl CO_2_-independent medium. The medium will preserve the correct pH value during the sorting procedure. Sorted cells can be centrifuged and re-suspended in normal cell culture medium prior to seeding in a proper cell culture dish (depending on the number of cells sorted). However, centrifugation is not recommended when total number of cells is very low (< 10,000).
7. If the integration of the AID-mRFP tag is very poor or not successful we recommend to use a recombinant adeno-associated virus system (rAAV) to deliver the repair template as described previously [21]. In this case, the efficiency of HDR is expected to be higher since the repair template is delivered as a linear ssDNA instead of a circular dsDNA plasmid.
8. If a single cell sort resulted in poor survival rates we recommend to sort 3 cells in a 96-well plate instead of 1 cell per well. After selection of a 3-cell population a limiting dilution can be performed to obtain a single clone. This will increase the chances of obtaining a positive clone.
9. At this step it is expected that at least one allele of the protein of interest (POI) is tagged. To achieve double heterozygous tagging more clones will likely need to be screened. Alternatively, a double selection strategy may be used (e.g. consecutive mRFP and EYFP tagging). If the protein is not tagged, we recommend to design different CRISPR/Cas9 guides and/or change the donor plasmid as described in Note 6, or to add to the construct a selectable cassette fused to a splicing site (e.g. P2A/T2A) or to a separate promoter. This latter strategy might be necessary if the POI is expressed at low levels and therefore not detectable by FACS. In the case of correct tagging but no protein depletion by IAA, a new target strategy needs to be generated. This might include moving the AID tag to a different protein extremity or adding a longer linker (in the case the AID is sterically hidden by the protein itself).
10. We designed primers that bind up- (forward) and down-stream (reverse) of the donor template or inside the AID sequence (reverse). HDR-mediated integration of the tag will result in a shift of the PCR product size when using AID-mRFP flanking primers (e.g. with tag integration: ˜1900 bp, without tag: ˜450 bp; Figure 1 C). A PCR product with the reverse primer binding inside the AID sequence is only expected in the presence of the AID tag.
11. Duration of auxin treatment is protein and cell line-dependent. Despite that for CENP-C we can observe complete degradation in only 20 minutes [11], for screening of correct AID integration we prefer to use an overnight treatment to be sure to achieve complete protein degradation.
12. Multiple washing steps are absolutely necessary to remove IAA properly. Washing needs to be done carefully since, if performed too harshly, it can detach cells from the dish.
13. Endogenous protein levels of CENP-C^AE^ and also CENP-B^mRFP^ are low in cells. This represents a problem to perform live-cell imaging since the RFP and EYFP signals are bleached over time. By acquiring images every 10 min we found a good compromise between total duration of acquisition (limited by bleaching) and time resolution to monitor CENP-C^AE^ reloading.
14. CENP-C^AE^ reloading cannot be observed immediately before mitosis (G2), during mitosis or early after mitosis (early G1) (Figure 2 B) as observed by filming asynchronous cells (the phases of the cell cycle were estimated by counting the time that the cells need to reach mitosis). However, we have found that CENP-C^AE^ is reloaded in all S-phase arrested cells (Figure 2 C). This is in agreement with our previously reported findings that CENP-C can only be loaded at the centromere in mid G1-phase (only after CENP-A deposition that occurs immediately at mitotic exit [22]) and in S-phase [8]. We have found that CENP-C levels re-accumulate linearly at the centromere after about 30-60 min (Figure 2 D), however we started to acquire images approximately 1h after IAA wash-out. Hence CENP-C^AE^ re-expression and reloading required a minimal time of about 90-120 min.

## ACKNOWLEDGEMENTS

The authors would like to thank all the members of the D. Fachinetti laboratory and C. Bartocci (I. Curie) for helpful suggestions. We thank Solène Hervé for providing images for figure 1 E. We also thank the FACS facility and the microscopy platform at Institut Curie. D.F. receives salary support from the CNRS. D.F. has received support by Labex « CelTisPhyBio », the Institut Curie and the ATIP-Avenir 2015 program. SH has received funding from the European Union’s Horizon 2020 research and innovation programme under the Marie Skłodowska-Curie grant agreement No 666003. This work has also received support under the program «Investissements d’Avenir » launched by the French Government and implemented by ANR with the references ANR-10-LABX-0038 and ANR-10-IDEX-0001-02 PSL.

